# Akt kinases are required for efficient feeding by macropinocytosis in *Dictyostelium*

**DOI:** 10.1101/417923

**Authors:** Thomas D. Williams, Sew-Yeu Peak-Chew, Peggy Paschke, Robert R. Kay

## Abstract

Macropinocytosis is an actin-driven process of large-scale, non-specific fluid uptake used for feeding by some cancer cells and the macropinocytosis model organism *Dictyostelium* discoideum. In *Dictyostelium*, macropinocytic cups are organised by ‘macropinocytic patches’ in the plasma membrane. These contain activated Ras, Rac and PI(3,4,5)P3 and direct actin polymerisation to their periphery. Here, we show that a classical (PkbA) and a variant (PkbR1) Akt protein kinase acting downstream of PI(3,4,5)P3 are together are near-essential for fluid uptake. This pathway enables the formation of larger macropinocytic patches and macropinosomes, thereby dramatically increasing fluid uptake. Akt targets identified by phosphoproteomics were highly enriched in small G-protein regulators, including the RhoGAP GacG. GacG knockout mutants make few macropinosomes but instead redeploy their cytoskeleton from macropinocytosis to motility, moving rapidly but taking up little fluid. The function of Akt in cell feeding through control of macropinosome size has implications for cancer cell biology.

**Summary statement:** *Dictyostelium* amoebae feed by macropinocytosis in a PIP3 dependent manner. In the absence of PI3-kinases or the downstream Akt protein kinases, cells have smaller macropinosomes and nearly abolished fluid uptake.

## Introduction

Macropinocytosis is an ancient process used by cells to take up large amounts of fluid (King and Kay, in press; Swanson, 2008). Actin-driven protrusions are extended in a cup-shape and then close to engulf extracellular medium into an internal vesicle (Buckley and King, 2017). Macropinocytosis occurs in a variety of mammalian cell types and clinically important settings, perhaps most notably by certain cancer cells as a way of obtaining nutrients (Bloomfield and Kay, 2016; Commisso et al., 2013; Kim et al., 2018; Palm et al., 2017).

Macropinocytic, and phagocytic, cups are formed by a ring of protrusive F-actin under the plasma membrane that is distinct from other large F-actin structures, such as pseudopods. Several cellular components involved in the organisation of these cups have been identified in mammalian and *Dictyostelium* cells, the most important of which seem to be Ras, Rac1 and the phospholipid PIP3 (and, by extension, its regulators PI3-kinases and PTEN) (Araki et al., 2007; Bar-Sagi and Feramisco, 1986; Fujii et al., 2013; Hoeller et al., 2013; Ridley et al., 1992; Veltman et al., 2016; Yoshida et al., 2009).

The proteins organizing macropinocytic cups are better known as members of both the growth factor signalling cascade and as oncogenes or tumour suppressors. Growth factors signal through receptor tyrosine kinases (RTKs), which activate downstream effectors including Ras (Margolis and Skolnik, 1994). Active Ras in turn activates class 1 PI3-kinases, recruited to the plasma membrane by RTKs, through their Ras binding domain to make PIP3, an interaction that is critical for growth of certain tumours (Castellano et al., 2013; Gupta et al., 2007; Hu et al., 1992). PI3-kinase activation leading to macropinocytosis can also occur independently of Ras (Palm et al., 2017). Activating mutations in the RTK/Ras signalling pathway occur in nearly half of cancers, and activating mutations of the PI3-kinase pathway, mostly PI3-kinases and PTEN, in a third, although these groupings include proteins not involved in macropinocytosis (Kandoth et al., 2013; Sanchez-Vega et al., 2018). Ras proteins are especially frequently mutated with, strikingly, ∼95% of pancreatic cancers driven by K-Ras (Fernandez-Medarde and Santos, 2011; Kandoth et al., 2013; Prior et al., 2012; Waddell et al., 2015). Further, loss of the RasGAP NF1, leads to increased Ras activation and tumour development (Bollag et al., 1996; Gutmann et al., 2017).

Macropinocytosis in the model organism *Dictyostelium discoideum* is highly analogous to macropinocytosis in mammalian cells, however the situation is simplified as growth factor signalling and RTKs are not present. In *Dictyostelium* macropinocytosis is used for nutrient acquisition and accounts for the vast majority of fluid uptake, making accurate quantification simple (Hacker et al., 1997; Williams and Kay, 2018a). Additionally, both forward and reverse genetic approaches can be employed (Bloomfield et al., 2015; Paschke et al., 2018).

The *Dictyostelium* macropinocytic cup is formed around a template ‘macropinocytic patch’ composed of activated Ras and Rac, PIP3 (although this is chemically different to mammalian PIP3) and F-actin, with a rim of the Arp2/3 activators SCAR/WAVE and WASP (Clark et al., 2014; Hoeller et al., 2013; Veltman et al., 2016). Other known components are Coronin, the myosin-I proteins and certain formins (Brzeska et al., 2016; Hacker et al., 1997; Junemann et al., 2016). The PIP3-phosphatase PTEN is excluded from the macropinocytic patch but is present on the rest of the plasma membrane (Hoeller et al., 2013; Iijima and Devreotes, 2002).

PIP3 is vital for efficient macropinocytosis, in both amoebae and mammalian cells (Araki et al., 1996; Hoeller et al., 2013; Williams and Kay, 2018b). However, despite its importance, its function in macropinocytosis is unknown. PIP3 acts by recruiting PIP3-binding proteins to the plasma membrane, often through a PH-domain. A considerable number of these proteins exist, but which are important for macropinocytosis is not known (Park et al., 2008; Zhang et al., 2010). The protein kinase Akt is an oncoprotein that is a major downstream effector of PIP3 in growth factor signalling and mTORC1 activation (Dibble and Cantley, 2015; Staal et al., 1977). Continuing the analogy with growth factor signalling, we examine its role in *Dictyostelium* macropinocytosis.

*Dictyostelium* has a classical Akt homologue called PkbA with a PIP3-binding PH domain that recruits it to the plasma membrane (Meili et al., 1999; Tanaka et al., 1999). It phosphorylates target proteins at the Akt consensus sequence and the phosphorylated proteins can be recognized with standard anti-phosphopeptide antibodies (Kamimura et al., 2008; Liao et al., 2010). In addition, there is variant enzyme, PkbR1, which is constitutively targeted to the plasma membrane by lipid modification and phosphorylates an overlapping set of target proteins (Kamimura et al., 2008; Liao et al., 2010; Meili et al., 2000). PkbA and PkbR1 have a conserved activation mechanism, requiring phosphorylation by TORC2 and PDK1 protein kinases (Alessi et al., 1997; Jacinto et al., 2006; Kamimura and Devreotes, 2010; Kamimura et al., 2008; Liao et al., 2010; Sarbassov et al., 2005; Stephens et al., 1998; Stokoe et al., 1997).

We show here that the Akt protein kinases, and their activating protein kinases, are required for efficient macropinocytosis. They act downstream of PIP3 to increase macropinosome size, and hence fluid uptake, by increasing the macropinocytic patch size. We identify some of the Akt targets by phosphoproteomics. Among these is the RhoGAP GacG, which is nearly essential for fluid uptake and without which cells move rapidly and make far fewer macropinosomes.

## Results

### PkbA and PkbR1 are together required for efficient fluid uptake by macropinocytosis

Fluid uptake by a *Dictyostelium* mutant lacking all Ras-activated PI3-kinases (PI3K1-5-) is practically abolished in shaking culture (Hoeller et al., 2013). However, some mutant strains have a conditional fluid uptake defect in shaking culture, but not when attached to a surface (Novak and Titus, 1997). We therefore re-assessed fluid uptake by surface-attached PI3-kinase mutants using flow-cytometry to measure uptake of fluorescent dextran. Fluid uptake was nearly abolished in PI3K1-/2- and other multiple PI3K mutants, confirming the importance of PIP3 for macropinocytosis in our assay conditions (Figure 1A).

**Figure 1:**
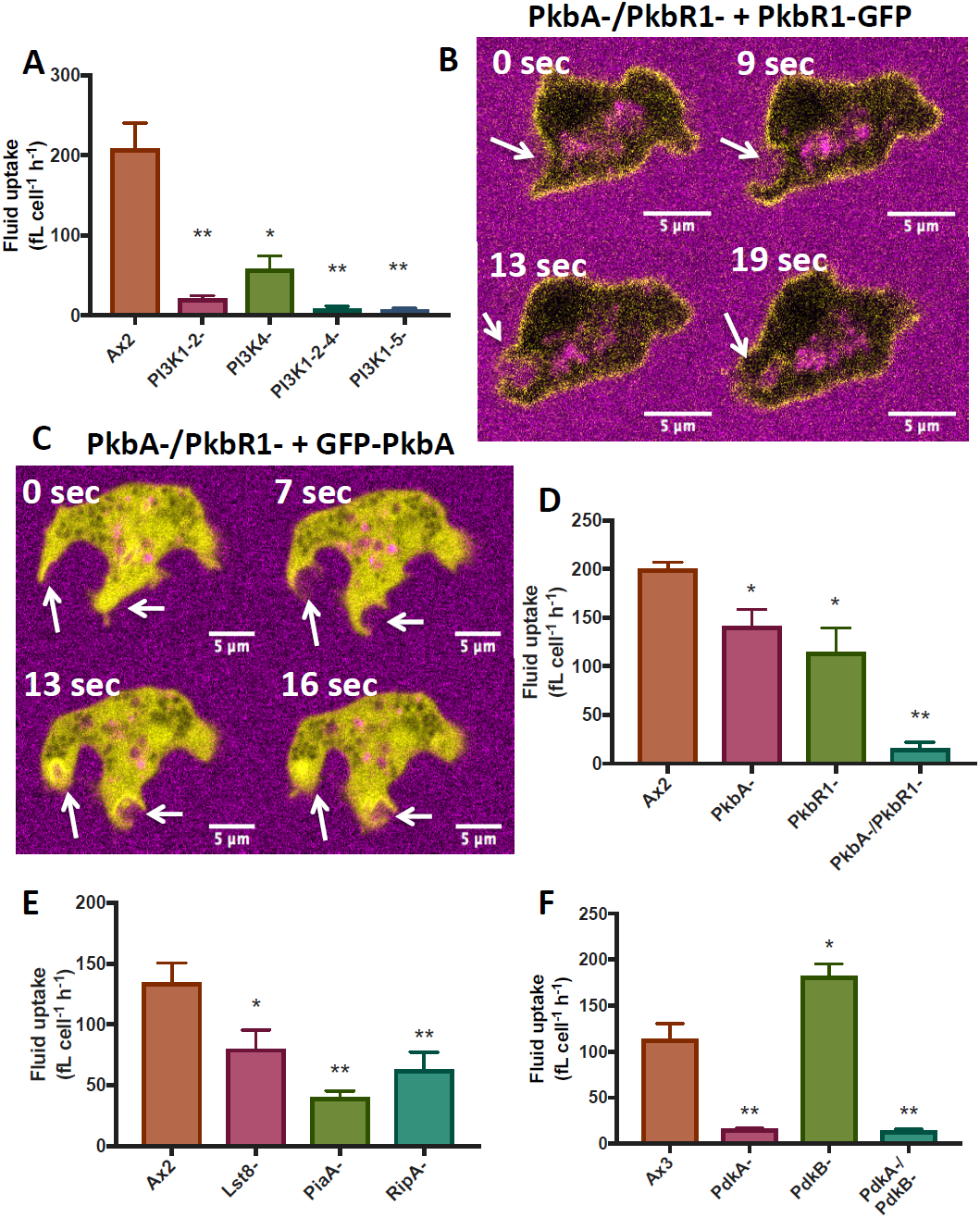
PkbA/PkbR1 as well as PI3-kinases are required for efficient macropinocytic fluid uptake. **A**) Fluid uptake is reduced in surface-attached PI3-kinase mutant strains, consistent with previous results in shaking suspension (Hoeller et al., 2013). **B**) PkbR1-GFP is localised on the plasma membrane of PkbA-/PkbR1-cells, including macropinosomes (arrow). TRITC-Dextran (magenta) marks the fluid-phase. **C**) GFP-PkbA is recruited to macropinosomes (arrows) in PkbA-/PkbR1-cells. TRITC-Dextran (magenta) marks the fluid-phase. **D**) Fluid uptake is practically abolished in a PkbA-/PkbR1-double mutant, similar to the PI3K1-5-strain. **E**) Fluid uptake is decreased in knockout mutants of components of the TORC2 complex, which activates PkbA/PkbR1 (n=6). **F**) Fluid uptake is nearly abolished in the absence of the PDK1 protein PdkA but increased by the absence of the alternative PDK1, PdkB. For fluid uptake experiments, cells were harvested from bacteria and inoculated in HL5 medium 24 h before measurement. Overexpression vectors were created using the pDM system (Veltman et al., 2009). Graphs show mean±SEM, n=3 unless specified. p≤0.1 is specified, * p<0.05, **p<0.01 compared to the parent strain.

The *Dictyostelium* Akt proteins, PkbA and PkbR1, may be downstream effectors of PIP3 in macropinocytosis. In vegetative cells, PkbR1 is present on the whole membrane, including macropinocytic cups (Figure 1B, Movie 1), while PkbA is recruited specifically to macropinocytic cups (Figure 1C, Movie 2), similar to Akt in mammalian cells (Yoshida et al., 2015). Akt could therefore be an evolutionarily conserved PIP3 effector during macropinocytosis.

Single and double knockouts of PkbA and PkbR1 (Figure S1A) were made in our Ax2 background using conditions that avoid the need for axenic growth (Paschke et al., 2018). Phosphorylation of Akt substrates was largely unaffected in the single knockouts and in the PI3K1-5-strain (Figure S1B) but was nearly abolished in the PkbA-/PkbR1-strain. This is likely explained by redundancy between PkbA and PkbR1, and PIP3 independent activation of PkbR1 in the absence of PIP3 (Kamimura et al., 2008) (Figure S1C).

Fluid uptake was modestly reduced in the PkbA- and PkbR1-single mutants, in agreement with previous data for PkbA-cells (Rupper et al., 2001). However, it was effectively abolished in the double mutant, showing that PkbA and PkbR1 are together essential for effective fluid uptake (Figure 1D), and phenocopying the PI3K1-5-strain.

Since PkbA and PkbR1 are together essential for efficient fluid uptake, we asked whether their activating kinases, TORC2 and PDK1 (Figure S1C) are also required. TORC2 has four subunits: Tor, PiaA/Rictor, Rip3/Sin1 and Lst8, all of which excluding Tor can be knocked out in *Dictyostelium*. These mutants have decreased, but not abolished, activation of PkbA and PkbR1 (Lee et al., 2005). In contrast to previously published results, fluid uptake was decreased in TORC2 component mutants (Figure 1E), although not to the same extent as in PkbA-/PkbR1-cells (Rosel et al., 2012).

*Dictyostelium* has two PDK1 homologs: PdkA and PdkB. PdkA, but not PdkB, is recruited to PIP3-containing membrane regions, such as macropinosomes, in a PI3-kinase dependent fashion, although binding to PIP3 has not been shown in-vitro (Kamimura and Devreotes, 2010; Liao et al., 2010). PdkA could therefore be responsible for PIP3-dependent activation of PkbA and PkbR1, like a classical PDK1 protein, with PdkB being responsible for PIP3-independent activation of PkbR1 (Figure S1C) (Alessi et al., 1997; Kamimura et al., 2008; Stokoe et al., 1997). In agreement with this model, fluid uptake by PdkA-mutants was effectively abolished, similar to PI3K1-5- and PkbA-/PkbR1-cells, but was actually increased in PdkB-cells (Figure 1F). PIP3-independent activation of PkbR1 by PdkB therefore has no function in macropinocytosis.

These results show that PkbA and PkbR1 and their activators, PdkA and TORC2, mediate efficient macropinocytic fluid uptake in *Dictyostelium*. The defect in PkbA-/PkbR1-cells is as severe as in PI3K1-5-cells and thus could account for their phenotype, though additional PIP3 effectors are not excluded.

### PkbA-/PkbR1-cells only have proliferation defects in axenic conditions

The fluid uptake defects observed in the PkbA-/PkbR1-cells could be due to a general defect in cell physiology. We therefore set out to investigate this.

PkbA-/PkbR1-cells do not proliferate in HL5 medium (Figure 2A), consistent with the large fluid uptake defect and a previously made mutant (Meili et al., 2000). PkbA-cells have a larger proliferation defect than their fluid uptake defect would indicate. Further investigation revealed that the rate of macropinocytosis decreases in these mutants on prolonged incubation in HL5 medium (Figures S2A&B), although the reason is unknown.

**Figure 2:**
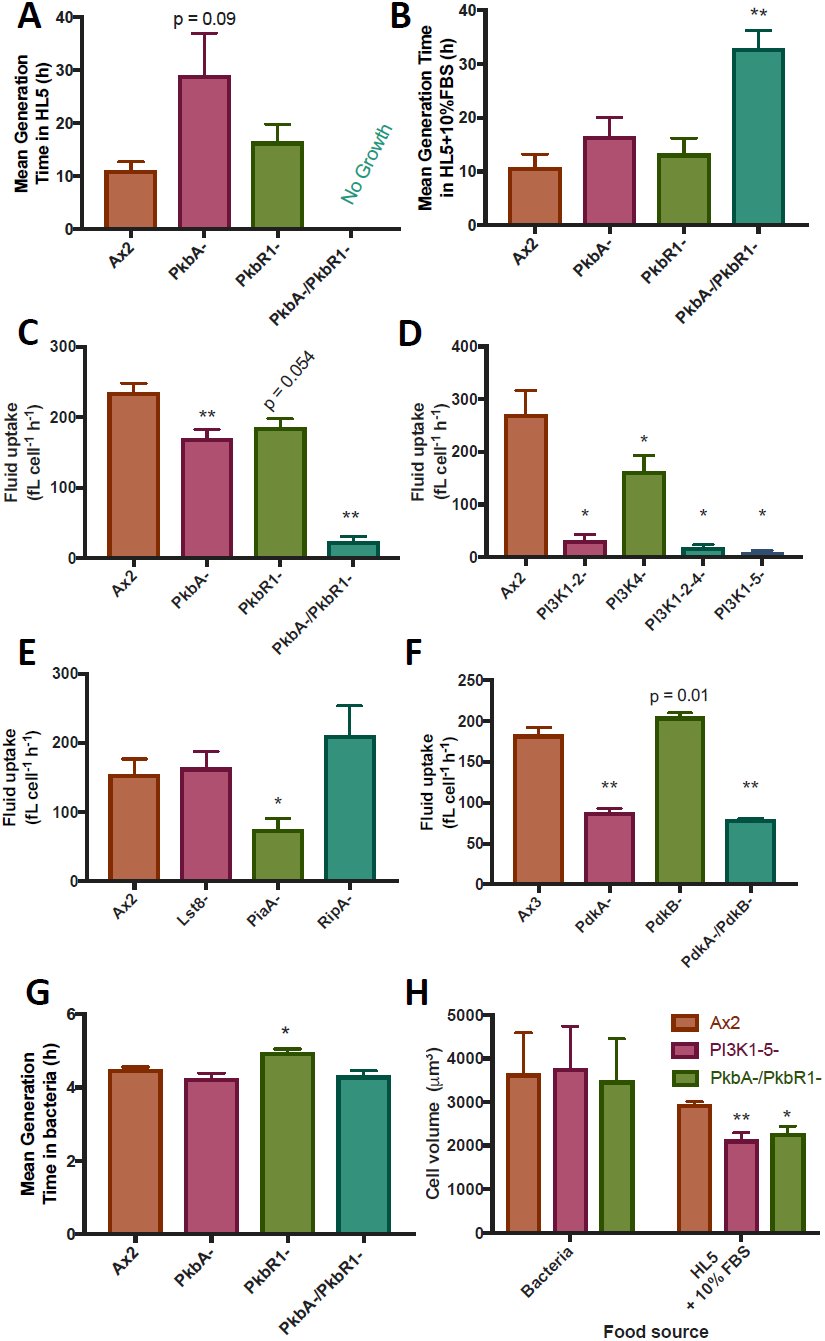
Cell proliferation and fluid uptake in enriched medium. **A**) PkbA-/PkbR1-cells do not proliferate in HL5 medium on a surface, while knockout mutants of just PkbA have a severe proliferation defect that does not correlate with their fluid uptake defect. Similar results were obtained in suspension (not shown) **B**) PkbA-/PkbR1-cells can proliferate when the HL5 medium is made more nutrient rich by addition of 10%FBS, albeit poorly. Proliferation of single mutants is also improved. Fluid uptake was therefore measured after 24 h incubation in HL5 medium + 10% FBS following cultivation on bacteria for: **C**) PkbA/PkbR1 mutants, **D**) PI3-kinase mutants, **E**) TORC2 component mutants, **F**) PDK1 mutants. PI3-kinase and PkbA/PkbR1 mutants had similar fluid uptake defects to when they were assessed in unsupplemented HL5 medium, while fluid uptake by TORC2 and PDK mutants was generally improved. **G**) Proliferation on bacteria is normal for all the PkbA/PkbR1 mutant strains. **H**) PI3K1-5- and PkbA-/PkbR1-cells are the same volume as Ax2 cells when cultivated on bacteria, where there is no proliferation defect (n=4). The mutant cell volume is relatively decreased after they have been incubated in HL5 medium + 10% FBS for 24 h (n=3), most likely due to the fluid uptake defect. Graphs show mean±SEM, n=3. p≤0.1 is specified, *p<0.05, **p<0.01 compared to Ax2.

The rate of macropinocytosis by Ax2 cells increases massively in the first 10 h after transfer to liquid medium, and this increase likely depends on nutrient sensing within the macropinocytic pathway (Williams and Kay, 2018b). If insufficient nutrition is obtained, upregulation does not occur. Poorly-macropinocytosing cells can sometimes grow and be stimulated into increased fluid uptake by medium enriched with 10% FBS (Bloomfield et al., 2015). Although growth and fluid uptake of the PkbA and PkbR1 single and double mutants could be improved by supplementation of the medium (Figure 2B), similar defects in fluid uptake were apparent (Figure 2C). This was also the case for the PI3-kinase mutants (Figure 2D). In contrast, the TORC2 component and PDK1 mutants, which can still partially activate PkbA/PkbR1 at macropinocytic cups, increase their fluid uptake in HL5 medium + 10% FBS, compared to HL5 only (Figures 2E&F). Further experiments using the PI3K1-5- and PkbA-/PkbR1-strains used HL5 medium + 10% FBS, where complications due to starvation are minimised. Further experiments using the TORC2 component and PdkA-mutants are performed in HL5 medium, as this supports their proliferation (Kamimura and Devreotes, 2010; Rosel et al., 2012).

In contrast to axenic growth, all PkbA/PkbR1 mutant strains proliferated rapidly on bacteria (Figure 2G), at similar rates to Ax2. PkbA-/PkbR1-, PI3K1-5- and Ax2 cells are a similar size when cultivated on bacteria, indicating the mutants have no abnormalities in cell cycle progression (Figure 2H). PI3K1-5- and PkbA-/PkbR1-cells are smaller than Ax2 when incubated in HL5 medium + 10% FBS, most likely due to the fluid uptake defect. Consistent with previous data, neither strain enters the developmental cycle when cultivated on bacterial lawns (Hoeller and Kay, 2007; Meili et al., 2000).

The normal growth of PkbA-/PkbR1-cells on bacteria and our inability to restore normal rates of fluid uptake to them under any nutritional condition tested indicate that they have a specific defect in macropinocytosis, rather than in general cell physiology or the regulation of macropinocytosis.

### Macropinocytic patches appear to be normally organized in PkbA-/PkbR1-cells

To investigate the cause of the fluid uptake defect in PkbA-/PkbR1-cells, we examined the organisation of their macropinocytic patches. Patches of the upstream components active Ras and PIP3 were still formed in the mutant (Figure 3A) and PTEN was excluded from them (Figure 3B), as expected. F-actin (Figure 3C) and activated Rac (Figure 3D) localisation, relative to active Ras, was also unperturbed in the mutant. SCAR/WAVE was examined using basal patches of active Ras, a surrogate for macropinosomes, as the marker is weak and easily bleached (Veltman et al., 2016) (Figure 3E). SCAR/WAVE was still enriched at the patch periphery. When this was quantified, no decrease in SCAR/WAVE enrichment at the ring was observed for the PkbA-/PkbR1-cells nor, in this instance, for the PI3K1-5-, contrary to expectations (Veltman et al., 2016).

**Figure 3:**
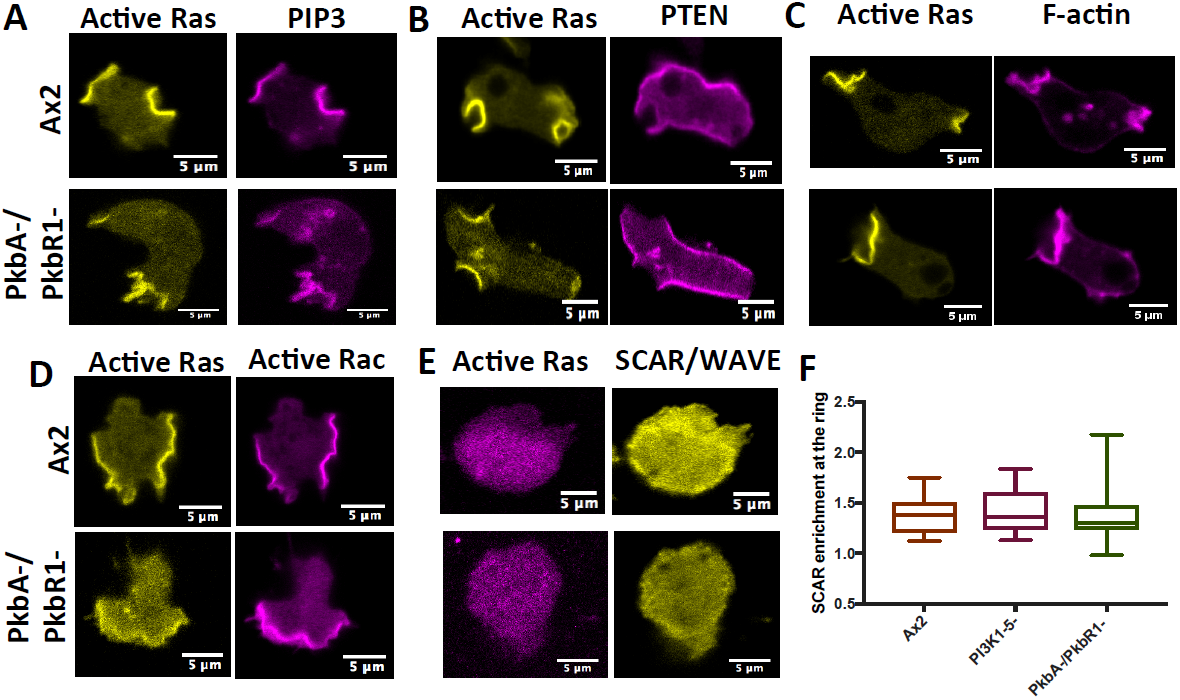
Macropinocytic patch components are normally localised in PkbA-/PkbR1-cells. Ax2 and PkbA-/PkbR1-cells were transformed with overexpression vectors containing markers of active Ras and another macropinocytic patch component. Representative cells show the relative localisation of each marker, which is not affected in the mutant. **A**) PIP3 (using PkgE-PH), **B**) PTEN (using PTEN), C) F-actin (using Lifeact), **D**) Active Rac (using PakB-CRIB), **E**) SCAR/WAVE (using HSPC300). HSPC300-GFP is technically challenging to visualise in macropinocytic patches, as it is faint and easily bleached. Basal patches were used as a surrogate (Veltman et al., 2016). **F**) The enrichment of SCAR/WAVE at the circumference of patches was no different between Ax2 and PI3K1-5- or PkbA-/PkbR1-cells, in contrast to previous data showing decreased recruitment in PI3K1-5-cells (Veltman et al., 2016). Cells were imaged on 3 separate days to obtain images of 32, 28 and 29 basal patches with rings of SCAR/WAVE in the 3 cell lines respectively. The SCAR/WAVE ring was defined in a Matlab script that measured the highest fluorescence along a set of normals following its rough outline (Veltman et al., 2016). This was divided by the SCAR/WAVE fluorescence inside the active Ras patch obtained using FIJI. Graph shows the minimum and maximum values, the box is the 25^th^ to 75^th^ percentile and the middle line shows the median.

### PkbA and PkbR1 increase the size of macropinocytic patches

We used microscopy to examine the rate at which macropinosomes form and their size. Macropinosomes form at a reduced rate in the PI3K1-5- and PkbA-/PkbR1-strains compared to Ax2 (Figure 4A), although this is not statistically significant and cannot account for the entire fluid uptake defect. We tested, and ruled out, the possibility that these macropinosomes were rapidly recycled to the extracellular medium by using a short uptake time (Figure S3A).

**Figure 4:**
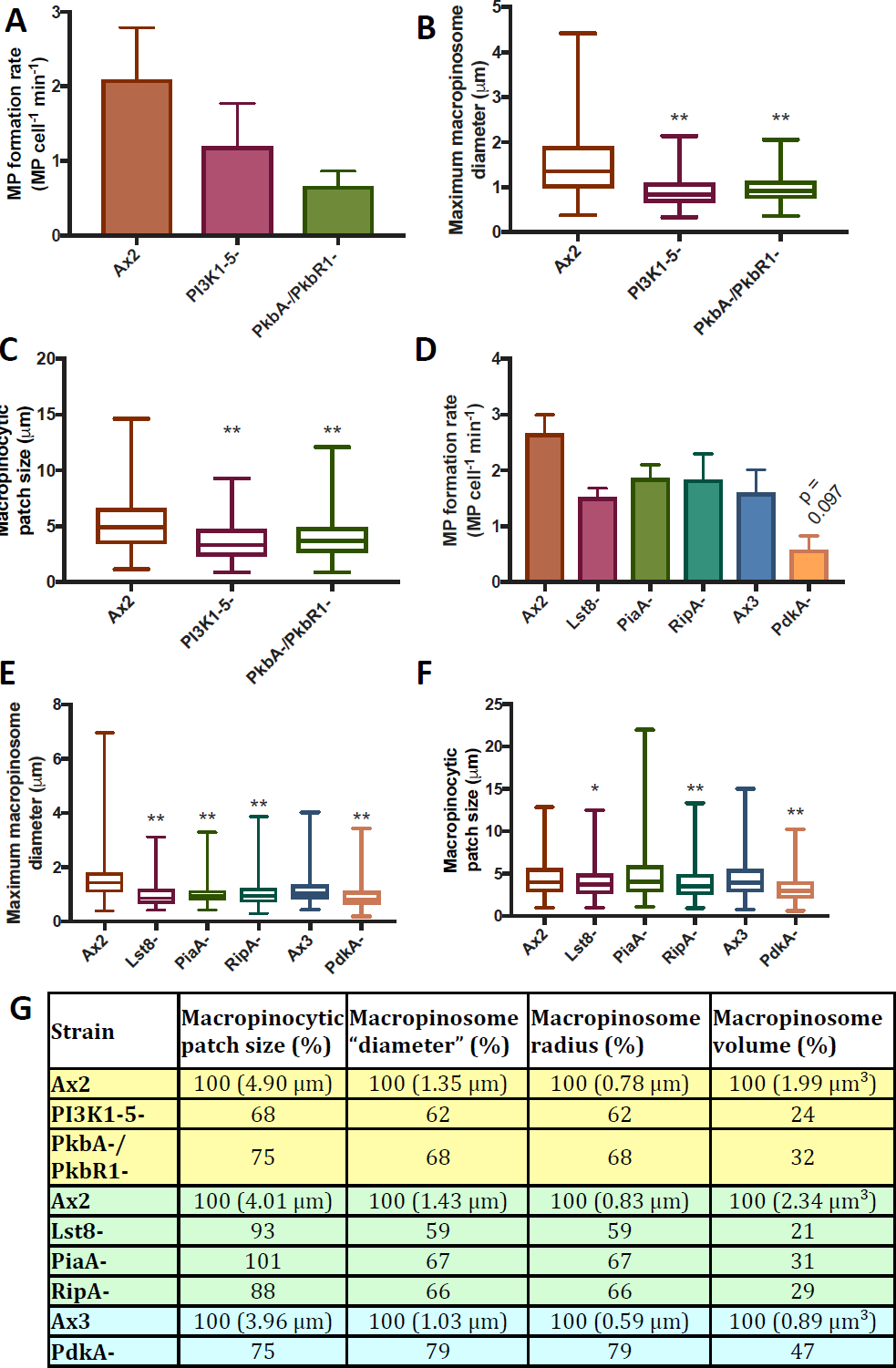
PkbA/PkbR1 regulate the size of macropinocytic patches. **A**) Macropinosome formation was reduced in both PI3K1-5- and PkbA-/PkbR1-strains compared to Ax2, but not by a statistically significant amount. **B**) Macropinosome diameter is decreased in PI3K1-5- and PkbA-/PkbR1-cells. This accounts for the majority of the fluid uptake defect. Macropinosomes analysed: Ax2 132, PI3K1-5-80, PkbA-/PkbR1-75. **C**) Macropinocytic patch size is decreased in PI3K1-5- and PkbA-/PkbR1-cells to a similar extent as the macropinosome diameter, indicating this is the reason for the macropinosome size decrease. Patches analysed: Ax2 453, PI3K1-5-168, PkbA-/PkbR1-330. **D**) Macropinosome formation is largely unaffected in TORC2 component mutants, but decreased for the PdkA-mutant. **E**) Macropinosome size is decreased in the TORC2 component and PdkA-mutants. Macropinosomes analysed: Ax2 233, Lst8-141, PiaA-176, RipA-211, Ax3 141, PdkA-202. **F**) Macropinocytic patch size is reduced in the PdkA-mutant, but is largely unaffected in the TORC2 component mutants. Patches analysed: Ax2 248, Lst8-289, PiaA-235, RipA-232, Ax3 307, PdkA-298. **G**) Summary table of the previous figures showing the average values as a percentage of the parent control. The measured diameter tends to halfway between the equator and the poles of the macropinosome (assuming the observed diameters were obtained from random sections of perfectly spherical macropinosomes). The average radius is calculated by multiplying the median observed diameter by tan(30°), and the volume calculated from that. As macropinosomes are often not perfect spheres when they are formed, and sampling at the periphery of macropinosomes is technically challenging, these values are approximate. Cells were harvested from bacteria and inoculated in medium (HL5 medium + 10% FBS for PI3K1-5- and PkbA-/PkbR1-, HL5 medium for PdkA- and TORC2 mutants, both with appropriate controls) for 24 h, then swapped to SUM before performing the experiment. Microscopy of macropinocytic patch and macropinosome size (maximum diameter as they closed) were performed on 3 separate days through random cross-sections of the cells, measured using FIJI and combined. Bar graphs show mean±SEM, box plots show the minimum and maximum values, the box is the 25^th^ to 75^th^ percentile and the middle line shows the median. n=3. p≤0.1 is specified, *p<0.05, **p<0.01 compared to the parent strain.

Instead, we find that PI3K1-5- and PkbA-/PkbR1-mutants form smaller macropinosomes. Macropinosome diameter at internalisation was reduced by ∼40% in both cases (Figure 4B), corresponding to a 3-4 fold decrease in macropinosome volume and thus accounting for a large proportion of the decrease in fluid uptake. A similar reduction in macropinocytic patch size was observed (Figure 4C), indicating a defect in macropinocytic patch size leads to smaller macropinosomes.

The size of particles that cells can phagocytose correlates with the size of macropinocytic patches formed (Bloomfield et al., 2015; Williams and Kay, 2018b). We measured phagocytosis of various size particles by PkbA-/PkbR1- and PI3K1-5-cells. The PkbA-/PkbR1- and PI3K1-5-mutants were less proficient than Ax2 at uptake of yeast (a comparatively large particle) (Figure S3C), while phagocytosis of smaller beads was unchanged (Figure S3D), consistent with the smaller macropinocytic patches made by these mutants.

The TORC2 component knockout and PdkA-mutants have similar phenotypes to the PI3K1-5- and PkbA-/PkbR1-cells. Macropinosome formation was reduced, but not to a statistically significant extent, in all the TORC2 component mutants, but was significantly reduced for PdkA-cells (Figure 4D). For all these mutants, macropinosome size was decreased (Figure 4E). However, while PdkA-mutants had a corresponding decrease in macropinocytic patch size, the TORC2 component mutants were largely unaffected in this regard (Figure 4F). This data is summarised in Figure 3G.

These results show that PI3K1-5- and PkbA-/PkbR1-cells make significantly smaller macropinocytic patches and macropinosomes than their parent, which (due to the cubic relation of linear dimensions and volume) leads to a much larger reduction in macropinosome volume. These results further imply that there is a local positive feedback loop in which the Akt protein kinases PkbA and PkbR1 increase the activity of upstream macropinocytosis components, thus enlarging macropinocytic patches.

### Identification of PkbA/PkbR1 targets by phosphoproteomics

The postulated feedback loop between upstream macropinocytosis components and PkbA/PkbR1 might operate indirectly and by many different potential routes, but ought to be mediated by targets of both PkbA and PkbR1. We therefore used an unbiased phosphoproteomic screen to identify such targets.

We compared the phosphoproteomes of parental Ax2 cells, PkbA-, PkbR1- and PkbA-/PkbR1-mutants and PkbR1-cells treated with LY294002 to inhibit PI3-kinase and hence PkbA. Phosphopeptides were isolated from these cells as described in the methods and the relative amounts of each phosphopeptide in each sample were compared by Tandem Mass Tag mass spectrometry over three biological replicates (Data S1). The phosphopeptides were filtered by reference to the fluid uptake data obtained earlier; ≥5-fold decreased in the PkbA-/PkbR1-mutant compared to Ax2, ≥2-fold decreased in the PkbR1-+ LY294002 cells with at least a 20% decrease in either of the PkbA- and PkbR1-samples. Only phosphopeptides appearing in at least 2 of the 3 repeats were accepted. Only 38 peptides from 30 proteins met these stringent criteria.

The kinases likely to be responsible for specific phosphorylations of the filtered set of peptides were predicted using NetPhos 3.1 (Blom et al., 2004). Proteins likely to be phosphorylated by PkbA/PkbR1 are shown in Table 1, while those more likely to be phosphorylated by other kinases are in Table S1. Of the potential PkbA/PkbR1 targets, half are predicted to function in signalling, while others are predicted to be involved in endosome trafficking and digestion (Figure 5A). GO analysis of the identified proteins revealed strong enrichment for terms associated with G-protein function, intracellular transport and enzyme function, while the most enriched term was for protein complex scaffold activity (Figure 5B). These predicted functions are consistent with proteins involved in cellular feeding by macropinocytosis. Additionally, several of these proteins have previously been identified as PkbA/PkbR1 targets, validating the approach (Charest et al., 2010; Kamimura et al., 2008; Tang et al., 2011).

**Table 1:** Candidate PkbA/PkbR1 effector proteins in macropinocytosis. Proteins predicted to be phosphorylated by PkbA/PkbR1 (Akt) that were less abundant in PkbA-/PkbR1-mutants (5-fold decreased in PkbA-/PkbR1-cells, 2-fold in PkbR1-+ LY294002 and a 20% reduction in either PkbA- or PkbR1-cells) compared to Ax2 after 24 h incubation in HL5 medium + 10% FBS to allow macropinocytosis upregulation and 30 min inhibitor treatment in KK_2_MC.

| Protein | Phosphorylation(s) | Probable kinase | Probable function | Closest Human homolog |
| --- | --- | --- | --- | --- |
| DDB_G0278629 | S675, S679 | GSK3, Akt | Unknown | LRRC16A |
| DDB_G0272206 | S1743 | Akt | Unknown | KIAA0664 |
| DDB_G0290337 | S483 | Akt | Unknown | N/A |
| DynC | S73 | Akt | Dynactin | Dynactin subunit 3 |
| GacG | S730 | Akt | RhoGAP | RhoGAP4 |
| GacH | S355 | Akt | RhoGAP | β-chimaerin 2 |
| GefS | S623 | Akt | RasGEF | SOS2 |
| GxcX | S1091 | Akt | RhoGEF | FGD6 |
| Kif6 | S1016 | Akt | Kinesin | KIF2C |
| KrsB | S505 | Akt | Protein Kinase | STK4 |
| LysA | S847, S851 | Akt, Cdc2 | Digestion | TLDC2 |
| ScaA | S225 | Akt | Scaffold protein | N/A |

**Figure 5:**
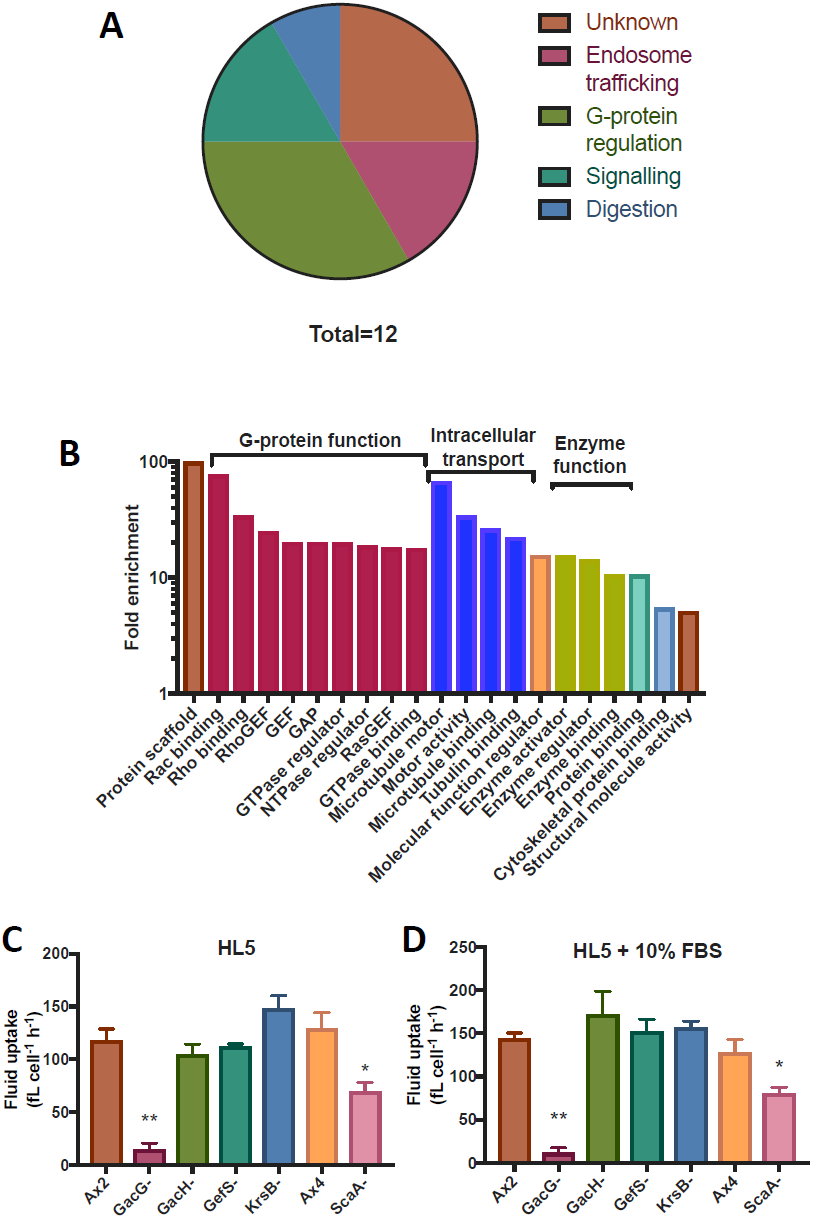
Fluid uptake by mutants of potential PkbA/PkbR1 targets. **A**) Probable functions of the identified PkbA/PkbR1 target proteins shown in Table 1. **B**) The identified proteins are highly enriched for ones involved in G-protein function, intracellular transport and enzyme function, while protein complex scaffold activity is the most enriched term. GO enrichment analysis was performed against the *D. discoideum* reference list using PANTHER to identify enriched molecular function terms with the Fishers exact test. Terms with a >5-fold enrichment are shown. **C**) Fluid uptake of the signalling mutants after 24 h incubation in HL5 medium after harvesting from bacteria. **D**) Fluid uptake of the signalling mutants after 24 h incubation in HL5 medium + 10% FBS after harvesting from bacteria. The fluid uptake of the mutants was similar between both media, with defects only seen for the GacG- and ScaA-mutants. Bar graphs show mean±SEM, n=3, *p<0.05, **p<0.01 compared to Ax2, ScaA-is compared to Ax4.

Since the PkbA/PkbR1-mediated increase in macropinocytic patch size most likely occurs through signalling events, we focused on the targets involved in G-protein function, protein complex scaffold activity and an identified protein kinase. A mutant of the ScaA scaffold protein was obtained from the *Dictyostelium* stock centre (Fey et al., 2013; Sawai et al., 2008), while knockout mutants were made of GacG, GacH, GefS and KrsB (Figure S4). We were unable to make a GxcX-mutant.

Fluid uptake of these mutants was measured in HL5 medium (Figure 5C) and HL5 medium +10% FBS (Figure 5D) and found to be decreased only in ScaA-cells (~50%) and GacG-cells (>90%, similar to PI3K1-5- and PkbA-/PkbR1-). That no defect was found for knockouts of the other proteins may be due to redundancy or compensatory elevated macropinosome formation rates that obscure a smaller macropinosome phenotype. As ScaA has previously been shown to be involved in a PkbA/PkbR1 activation feedback loop, we focused on GacG, which has a RhoGAP and a FERM domain (Charest et al., 2010).

### GacG-mutants have decreased macropinocytosis and increased motility

Loss of GacG, as a RhoGAP, is expected to lead to over-activation of one or more Rho proteins. Experiments in mammalian cells have shown that Rac1, a Rho protein, must be inactivated for macropinosome internalisation, and data suggests a similar function in *Dictyostelium* (Dumontier et al., 2000; Fujii et al., 2013). If Rac1 inactivation does not occur, arrested macropinocytic structures can accumulate. To test if such a scenario occurs in GacG-cells, we examined macropinocytic patches using live-cell confocal microscopy. No obvious defect in macropinosome internalisation was observed, and neither was there an accumulation of macropinocytic structures. In fact, there were far fewer macropinocytic patches formed (Figure 6A). The macropinosome formation rate is also greatly reduced (Figure 6B), and those that do form are smaller (Figure 6C). Despite the fluid uptake defect, GacG-cells obtain sufficient nutrients to proliferate in HL5 medium + 10% FBS, although much slower than Ax2 cells (Figure 6D). The comparative lack of macropinocytosis is therefore not due to the cells prematurely entering the developmental cycle when in axenic conditions.

**Figure 6:**
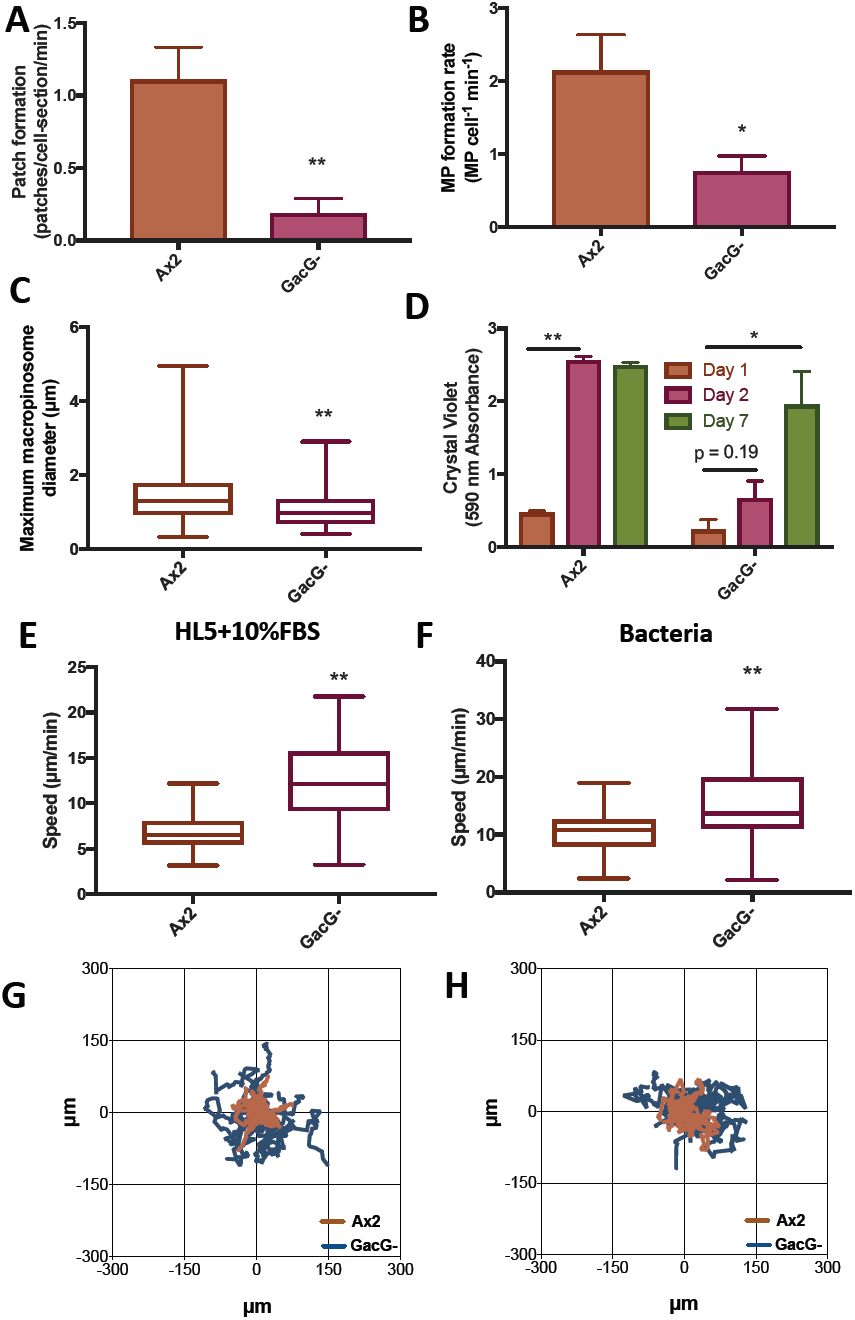
Macropinosome formation is severely impaired in GacG-cells but their motility increased. **A**) GacG-mutant cells make fewer macropinosomes than Ax2 cells (n=4). **B**) GacG-mutant cells make smaller macropinosomes than Ax2 cells. Cells expressing an active Ras marker were filmed on 7 separate days, macropinosomes measured at internalisation and the results combined. Macropinosome number: Ax2 254, GacG-63. **C**) GacG-cells make fewer macropinocytic patches, accounting for the decrease in macropinosome formation. Cells expressing a marker for active Ras were filmed through a random cross section for 5 min and the number of macropinocytic patches formed per cell-section per min was calculated (n=4). **D**) GacG-cells proliferate when incubated in HL5 medium + 10% FBS. Ax2 and GacG-cells were harvested from bacteria, 5×10^4^ inoculated into 0.5 ml HL5 medium + 10% FBS in triplicate in 24-well tissue culture plates and proliferation scored using crystal violet staining after 1, 2 or 7 days. Ax2 cells became confluent after 2 days, whereas this took much longer in the case of GacG-cells. **E**) GacG-cells move faster than Ax2 cells after incubation in HL5 medium + 10% FBS. **F**) GacG-cells move faster than Ax2 cells after incubation in bacterial suspension. G) Tracks of Ax2 and GacG-cells after incubation in HL5 medium + 10% FBS. H) Tracks of Ax2 and GacG-cells after incubation in bacterial suspension. Cells were harvested from bacteria and incubated in HL5 medium + 10% FBS for 24 h before performing the experiments unless otherwise specified. The speed of random movement of 120 cells for each strain was measured in E) and F) and and the tracks of 20 cells for 20 min shown in G) and H). Bar graphs show mean±SEM, box plots show the minimum and maximum values, the box is the 25^th^ to 75^th^ percentile and the middle line shows the median. n=3 unless specified. *p<0.05, **p<0.01 compared to Ax2.

The GacG-fluid uptake defect was not rescued by overexpression of either GacG-GFP or GAP-dead GFP-GacG, which both localise evenly throughout the cytosol (not shown), while cells were highly resistant to overexpression of GFP-GacG, hampering further investigation of GacG function and localisation.

Macropinocytosis and cell movement both require the construction of large actin-driven projections by cells – cups and pseudopods – and in active cells may be in competition for limited cytoskeletal resources (Vargas et al., 2016; Veltman et al., 2014). We noticed that GacG-cells moved much more freely and faster than the parent Ax2 strain. The difference is greatest with cells grown in both HL5 medium + 10% FBS (Figure 6E), where macropinocytosis is favoured in Ax2. This difference is still present, but is reduced, when cells are grown on bacteria, where Ax2 macropinocytosis is lower (Figure 6F). The differences are also clear in cell tracks (Figures 6G&H).

GacG-cells take up much less fluid than Ax2 cells due to a decrease in macropinocytic patch (and thereby macropinosome) formation and appear to have switched cytoskeletal resources from macropinocytosis to movement, moving considerably faster in all circumstances investigated.

## Discussion

In this work we show that, together, the Akt protein kinases PkbA and PkbR1 are nearly essential for fluid uptake in *Dictyostelium*. The defect in fluid uptake by PkbA-/PkbR1 cells is as great as that for cells lacking all Ras-activated PI3-kinases. In both mutants, this is because of a decrease in macropinosome volume due to a decrease in macropinocytic patch size. As Akt acts downstream of PIP3, lack of activation of Akt could account for the phenotype of the PI3-kinase mutants, although a role for other PIP3-binding proteins is not excluded.

PIP3 is required for effective macropinocytosis in mammalian cells, but the role of Akt is less clear (Araki et al., 1996). Inducing macropinosome formation activates Akt at macropinosomes, and this activation can be inhibited with macropinocytosis inhibitors (Chiasson-MacKenzie et al., 2018; Erami et al., 2017; Yoshida et al., 2018 preprint). However, macropinosome formation is not affected by inhibition of Akt in macrophages (Pacitto et al., 2017; Yoshida et al., 2015). In other cell-types, the opposite is true: Akt inhibition inhibits macropinocytic uptake of collagen by stellate cells (Bi et al., 2014). Akt1 deletion reduces tumour development in mice with activated Ras or PTEN deficiency, mutations which allow tumour cells to use macropinocytosis for nutrient acquisition (Chen et al., 2006; Commisso et al., 2013; Kim et al., 2018; Skeen et al., 2006). The functional relevance of Akt activation at macropinosomes remains to be fully investigated it could, in a subset of cell-types, increase macropinosome volume and fluid uptake.

Macropinocytic cups form spontaneously in the plasma membrane of *Dictyostelium* cells, without the need for receptor stimulation. Therefore, the role of the Akt protein kinases PkbA and PkbR1 in macropinocytosis is unlikely to be in conventional signal transduction to activate macropinocytosis. The evidence presented here shows that a major function of PkbA and PkbR1 is to make these cups larger, and so able to engulf more liquid, or in the case of phagocytosis, take up larger particles. This role, in which Akt activation leads to a local increase in activated Ras (as indicated by larger macropinocytic patches) and hence its activator, PIP3, suggests that Akt is part of a local positive feedback loop.

Our search by phosphoproteomics for targets of Akt and possible components of the postulated activating feedback loop yielded a small list of proteins. Among these we focussed on GacG, a putative RhoGAP, mutants of which have nearly abolished fluid uptake and increased movement. Macropinocytic cups and pseudopods are both substantial F-actin structures and may compete for the same cytoskeletal resources in active cells. In *Dictyostelium*, cells grown in liquid medium produce many macropinosomes but move poorly, while it is the reverse for cells grown on bacteria, as well as cells undergoing development (Fisher et al., 1989; Veltman et al., 2014; Williams and Kay, 2018b). This competition between macropinocytosis and movement is also observed in dendritic cells (Vargas et al., 2016). GacG-cells are phenotypically similar to cells grown on bacteria when growing in axenic conditions, indicating GacG may somehow be required for proper axenic adaptation. It is possible that GacG might act as a switch between pseudopods and macropinosomes by suppressing pseudopod formation.

Ras, the RasGAP NF1, Ras-activated PI3-kinase, the PIP3 phosphatase PTEN and Akt are all important for macropinocytosis in *Dictyostelium*. In mammalian cells, Ras, PI3-kinase, PTEN and, in some instances, Akt have likewise been linked to macropinocytosis. These proteins are better known as oncogenes or tumour suppressors. Both activated Ras and PTEN deficiency stimulate macropinocytosis in mammalian cells, supporting proliferation (Commisso et al., 2013; Kim et al., 2018; Palm et al., 2017). PI3-kinase inhibition inhibits macropinocytosis and inhibition of Akt may have a similar effect in certain cell-types (Araki et al., 1996). It is possible that activating mutations in PI3-kinases and Akt may similarly support increased nutrient acquisition by macropinocytosis. If this is so, inhibiting macropinocytosis may be an effective way of inhibiting the growth of tumours activated in different components of the RTK-Ras-PI3-kinase-Akt pathway.

## Materials and Methods

### Dictyostelium *culture conditions*

*Dictyostelium* discoideum Ax2 cells and derivatives were used in this study (See supplementary table 1) and were cultivated on *Klebsiella aerogenes* bacteria on SM plates at 22°C unless otherwise stated. Nutrient media (HL5 (Formedium, UK), HL5 + 10% FBS and SUM (Williams and Kay, 2018b)) were supplemented with Dihydrostreptomycin (100 µg/ml), Ampicillin (100 µg/ml) and Kanamycin (50 µg/ml). Strains were stored frozen in Horse Serum + 7.5% DMSO in N_2_ (l) and scraped out using a 16G hypodermic needle (Becton Dickinson) onto *K. aerogenes* and SM agar when needed. Typically cells were passaged once a week for 4-5 weeks before being refreshed. Except where indicated, we used a standardized set of mutants created in our laboratory strain of Ax2, most of which were made using transformation conditions that did not depend on axenic proliferation, reducing the risk of suppressing mutations.

### Cell uptake measurements

All cell uptake measurements were performed as described (Williams and Kay, 2018a; Williams and Kay, 2018b). Briefly, cells were cultivated on SM agar with *K. aerogenes* bacteria, harvested, washed free of bacteria and inoculated in the indicated nutrient medium to 1×10^5^ cells/ml, with triplicate 50 µl samples in wells of a 96-well plate. This was incubated for 24 h at 22°C before 50 µl of the same medium containing 1 mg/ml TRITC-Dextran (155 kDa, Sigma-Aldrich) was added for 1 h unless otherwise specified. This was then thrown off, the plate washed by submersion in ice-cold KK_2_ buffer (16.6 mM KH_2_PO_4_, 3.8 mM K_2_HPO_4_, pH 6.1) and the cell detached using 100 µl 5 mM sodium azide dissolved in KK_2_MC (KK_2_ + 2 mM MgSO_4_, 100 µM CaCl_2_). The fluorescence of each cell in the samples was then measured by an LSR_II flow cytometer (BD Biosciences) and analysed using FlowJo.

For measurement of phagocytosis, cells were harvested from bacteria as above. In the case of 1.5 µm bead (Polysciences) uptake, the cells were diluted into KK_2_MC and plated as before, and given ∼30 min to settle. 1.5 µm YG beads were washed free of sodium azide and diluted into KK_2_MC at 2×10^8^ beads/ml. 50 µl of this was added to each well for 20 min, after which cells were washed, detached and internalised fluorescence measured as before and analysed as described (Sattler et al., 2013). For yeast uptake, the cells were resuspended in 5 ml conical flasks at 5×10^6^ cells/ml and shaken for ∼30 min before addition of 1×10^7^ sonicated TRITC-yeast particles/ml. 200 µl samples were taken at 0 and 60 min, mixed with 2 µl 0.4% Trypan Blue (Sigma-Aldrich) by shaking for 3 min, washed twice in KK_2_ + 10 mM EDTA and the fluorescence determined on a Perkin Elmer LS 50 B fluorimeter (excitation at 544 nm, emission at 574 nm, each with a 10 nm slit).

### Transformation of Dictyostelium

Vectors were made according to standard procedures, propagated using *Escherichia coli* XL10 and harvested using the ZR miniprep classic kit (Zymogen).

Approximately 1×10^6^ *Dictyostelium* cells were harvested from growth zones on bacterial plates into 1 ml H40 buffer, pelleted by centrifugation at max speed on a benchtop centrifuge for 10 sec and resuspended in 100 µl H40 (Paschke et al., 2018). Vector was added to the cells (500 ng overexpression vector, 2 µg linearised knockout vector) and they were chilled on ice with 2 mm electroporation cuvettes (SLS). The cells were transferred to the 2 mm cuvettes then subject to square wave electroporation in a Bio-Rad GenePulser Xcell (2 × 350 Volts for 8 ms, with a 1 sec gap) and added to 2 ml KK_2_MC + 2 OD_600 nm_ *K. aerogenes* in a 3.5 cm dish to recover for 5 h.

After recovery, for cells transformed with an overexpression vector the appropriate selection (10 µg/ml G418, although this was doubled when working with the PkbA-/PkbR1-cells due to poor marker expression, and 100 µg/ml hygromycin) was added to the dish and the dish swirled to ensure an even distribution. In the case of knockout transformations, two 50 ml falcon tubes containing 30 ml KK_2_MC + 2 OD_600 nm_ *K. aerogenes* and selection were prepared, and the dish split between the two of them (200 µl and 1.8 ml). These were vortexed to mix and 150 µl put into each well of two 96-well plates per tube, and incubated at 22 °C. Transformants were typically obtained after ∼4 days (overexpression) and ∼6-8 days (knockout).

### Screening for mutants

Confluent wells of cells resistant to drug were obtained. To screen these, the media in the well was pipetted up and down to resuspend the cells and 2 µl from the well was added to 20 µl lysis buffer with 20 µg/ml freshly added proteinase K in a 0.2 ml PCR tube (Paschke et al., 2018). The tube was vortexed to mix, then the proteinase K inactivated at 95°C for 1 min. 2 µl of DNA was added to a 25 µl PCR reaction to screen from the resistance cassette to outside the construct, giving a product that could only derive from a mutant.

Once wells containing mutants were identified, these were plated clonally onto SM plates with *K. aerogenes* and individual colonies were screened (colony appearance usually took 4 days), with DNA this time being prepared using a g-DNA miniprep kit (Zymogen).

To make double mutants, the resistance cassette in a single mutant strain was removed by Cre-Lox recombination: 500 ng of pDM1489 was transformed into cells, selected for, and then plated out on SM plates with *K. aerogenes* to remove the selective pressure. Clonal populations that had lost both resistance markers were identified; sometimes this took more than one passage of cells on bacteria.

### Microscopy

#### Macropinosome formation rate

Cells were harvested from bacterial plates, washed free of bacteria and incubated in 2 ml of the indicated medium for 24 h at 1×10^5^ cells per well of a 6 well plate. The medium was removed and the cells resuspended in SUM and transferred to a 2-well microscope slide (Nunc) and allowed to settle for ∼ 1 h. The medium was removed and 0.5 ml SUM containing 2 mg/ml FITC-dextran (70 kDa, Sigma-Aldrich) was added for 1 min, removed and the cells washed 2x with KK_2_MC, fixed using 4% paraformaldehyde for 20 min, and washed 5x with PBS (pH 5) and stored at 4°C. Z-stacks with 0.1 µm steps were taken using a Zeiss 700 series microscope and the number of dextran positive vesicles counted manually using FIJI.

#### Live cell microscopy

Cells were transformed with overexpression vectors as above. Transformants were washed free of bacteria and 1×10^5^ were incubated in 2 ml medium for 24 h in a 6 well plate at 22°C. The medium was removed and replaced with SUM, and the cells transferred to 2-well microscope slides for ∼30 min before imaging using a Zeiss 700 series microscope. Movies were typically taken for 5 min with 1 frame per sec. The active Ras marker Raf1-RBD was used to observe macropinosomes and macropinocytic patches unless otherwise specified. When measuring macropinosome diameters, random cross-sections of cells were used meaning any individual diameter measured using this technique could be at any section through the macropinosome, but in the aggregate the average tends to halfway between the equator and the poles.

#### Motility assay

Cells were cultivated on SM agar plates in co-culture with *Klebsiella aerogenes* bacteria. 24 h before the assay, cells were transferred into 2 ml of either HL5 medium + 10% FBS or a *K. aerogenes*/KK_2_MC suspension (OD_600 nm_ = 2) at 2×10^5^ cells/ml and plated into 6-well dishes. After 24 h, cells were washed 5 times in KK_2_MC and resuspended in the same buffer to a final density of 1×10^5^ cells per ml. The cells were finally seeded in a MatTek 35 mm petri dishes with a 20 mm 1.5 coverglass and allowed to attach for 20 min. The random motility of the amoebae was observed for 30 min, taking frames every 30 sec using a Zeiss 700 series microscope equipped with a 10x air objective. Automatic cell tracking was performed as described previously (Susanto et al., 2017).

### Mean Generation Time determination

To measure proliferation in shaking axenic culture, cells were cultured to logarithmic phase in shaking axenic culture, diluted to 5×10^5^ cells/ml in fresh medium and counted approximately every 12 h using a haemocytometer.

To measure axenic proliferation of surface attached cells; cells were prepared as above, then 2×10^4^ were plated in 0.5 ml fresh medium in duplicate in multiple 24 well plates which were placed in a 22°C incubator. Approximately every 12 h, a plate was removed from the incubator and the media aspirated. The wells were washed once with buffer before staining with 0.1% crystal violet dissolved in 10% ethanol for 20 min. The free crystal violet was removed by washing three times with water, after which the crystal violet staining the cells was dissolved by addition of 0.9 ml 10% acetic acid for 20 min. The absorbance at 590 nm of the resulting solution was obtained using a Nano Drop 2000 (Thermo Scientific) and the background crystal violet staining from a set of wells with no cells was subtracted.

Finally, to measure proliferation with bacteria as the food source, cells were cultivated on SM plates in conjunction with *K. aerogenes*. 1×10^4^ cells/ml from growth zones were seeded in a conical flask containing 5 ml KK_2_MC + K. aerogenes (at 20 OD_600 nm_) and the cell density counted approximately every 4 h during the daytime using a haemocytometer.

### Western Blots

Cells were harvested from SM plates and washed free of bacteria. 1×10^6^ cells were inoculated into 10 ml HL5 medium + 10% FBS in a 9 cm tissue culture plate for 24 h. They were then resuspended by pipetting, pelleted by centrifugation at 370 x *g* for 3 min, the supernatant discarded, and washed twice in 50 ml ice-cold KK_2_MC buffer. Cells were counted and transferred to a 1.5 ml eppendorf tube, spun down at max speed in a benchtop centrifuge for 10 s, the supernatant removed by aspiration, resuspended to 2×10^7^ cells/ml in 1x sample buffer (NuPage) containing 2.5% 2-mercaptoethanol and protease and phosphatase inhibitors and incubated for 5 min at 95°C.

10 µl of the cell solutions were run out on 4-12% Bis Tris 10 well gels (Nupage) with See Blue Plus 2 used as a ladder. These were blotted onto immobilon P (Merck) membranes. This was blocked with 2% BSA (Fisher Scientific) in TBS-T for 2-3 h, then incubated with primary antibody (DM1A and 23C8D2, CST) overnight in 5% BSA in TBS-T. Membranes were rinsed twice with TBS-T and washed thrice for 10 min before addition of the secondary antibody conjugated to HRP (172-011, BioRad and Ab97051, Abcam) in 5% BSA for 2-3 h, then washed again. Antibodies were imaged with GE Healthcare detection reagent on a BioRad chemidoc using the chemidoc hi-resolution setting. Blots were stripped using restore PLUS western blot stripping buffer (Thermo-fisher) as per the manufacturers instructions before re-probing.

### Cell Size

Cell size was determined using an Eclipse flow cytometer (Sony iCyt). Cells were taken directly from their growth condition/treatment and, without washing, passed through a 70 µm filter (Sysmex CellTrics), and run through the Eclipse, which measured the median diameter and volume of the cell population.

### Mass Spectrometry

#### Sample preparation for mass spectrometry

Approximately 1-2×10^8^ vegetative cells for each strain (Ax2, PkbA-, PkbR1-, PkbA-/PkbR1-) were harvested from cultivation on bacterial SM agar plates and washed five times in KK_2_MC using benchtop centrifugation at 280 x *g* for 3 min. The harvested cells were resuspended to 2×10^6^ cells per ml in HL5 medium + 10% FBS, and incubated in a shaking conical flask at 22° C for 24 h.

The cells were then pelleted in a benchtop centrifuge as before, washed twice in KK_2_MC and resuspended to 1×10^7^ cells per ml in KK_2_MC. The PkbR1-cells were split between two conical flasks, one of which had LY294002 (Cayman Chemical) added to 100 µM (to prevent PkbA recruitment to macropinosomes and activation) while the rest of the samples all had 0.2% DMSO vehicle added (Ax2 + DMSO = sample 1, PkbA-+ DMSO = sample 2, PkbR1-+ DMSO = sample 3, PkbR1-+ LY294002 = sample 4, PkbA-/PkbR1-+ DMSO = sample 5). The cells were incubated shaking at 22 °C, 180 rpm for 30 min, then mixed in a 1:1 ratio with 10% TCA, and incubated on ice for at least 30 min to lyse the cells and precipitate the protein.

The protein was pelleted in a benchtop centrifuge by spinning at 2400 x *g* for 10 min, then washed twice with 20 ml of ice-cold acetone under the same conditions to remove the TCA. The protein was resuspended in protein solubilisation buffer (8M Urea, 20 mM HEPES, pH 8) and the concentration of the samples was measured using a Bradford assay (BioRad) and adjusted to 2.5 mg/ml, frozen in dry ice and stored at −80° C.

#### Enzymatic Digestion

Following the isolation of protein, 450 µg of each sample was reduced with 5 mM DTT at 56°C for 30 min then alkylated with 10 mM iodoacetamide in the dark at room temperature for 30 min. They were then digested with mass spectrometry grade Lys-C (Promega) at a protein: Lys-C ratio of 150: 1 (w/w) for 4 h at 25°C. Next, the samples were diluted from 8 M to 1.5 M urea using 20 mM HEPES (pH 8.5) and digested at 30°C overnight with trypsin (Promega) at a 75: 1 (w/w) protein: trypsin ratio. Digestion was stopped by the addition of trifluoroacetic acid (TFA) to a final concentration of 1%. Any precipitates were removed by centrifugation at 13000 x *g* for 15 min. The supernatants were desalted using homemade C18 stage tips containing 3M Empore extraction disks (Sigma-Aldrich) and 8 mg of poros R3 (Applied Biosystems) resin. Bound peptides were eluted with 30-80% acetonitrile (MeCN) in 0.1% TFA and lyophilized.

#### TMT peptide labelling

The lyophilized peptides from each sample were resuspended in 75 µl of 3% MeCN. The peptide concentrations were determined by Pierce Quantitative Colorimetric Peptide assay (Thermo Scientific) according to the manufacturers’ instructions, except the absorbance was measured by Nanodrop Spectrophotometers (Thermo Scientific) at 480 nm. 0.8 mg of each TMT 10plex reagent (Thermo Scientific) was reconstituted in 41 µl anhydrous MeCN. The peptides from each of the samples were labelled with a distinct TMT tag in 170 mM triethylammonium bicarbonate for 1 h at room temperature (r.t.). The labelling reaction was stopped by incubation with 8 µl 5% hydroxylamine for 15 min. The labeled peptides from each repeat were combined into a single sample and partially dried to remove MeCN in a SpeedVac (Thermo Scientific). After this, the sample was desalted as before and the eluted peptides were lyophilized.

#### Phosphopeptide enrichment

The lyophilized labeled peptides were resuspended in 1.2 ml of 50% MeCN, 2 M lactic acid (loading buffer) in an Eppendorf tube and incubated with 20 mg TiO_2_ beads (Titansphere, GL Sciences, Japan) at room temperature for 1 h. For the second round of enrichment, the beads slurry was spun down by centrifugation, and the supernatant was transferred to another Eppendorf tube. The supernatant was incubated with 15 mg of TiO_2_ beads and the beads slurry was spun down again as before. The beads were transferred to homemade C18 stage tips, washed in the tip twice with the loading buffer and once with 50% MeCN, 0.1% TFA. Phosphopeptides were eluted sequentially with 50 mM K_2_HPO_4_ (pH 10) followed by 50 mM K_2_HPO_4_, 50% MeCN (pH 10) and 50% MeCN, 0.1% TFA. The eluates were partially dried in a SpeedVac after acidification and were desalted as described above.

#### Basic pH Reverse-Phase HPLC fractionation

The enriched phosphopeptides were subject to off-line high performance liquid chromatography (HPLC) fractionation, using an XBridge BEH130 C18, 5 µm, 2.1 mm x 150 mm (Waters) column with an XBridge BEH C18 5 µm Van Guard cartridge, connected to an Ultimate 3000 Nano/Capillary LC System (Dionex). Phosphoeptides were resolubilised in solvent A (5% MeCN, 95% 10 mM ammonium bicarbonate, pH 8) and separated with a gradient of 1-90% solvent B (90% MeCN, 10% 10 mM ammonium bicarbonate, pH 8) over 60 min at a flow rate of 250 µl/min. A total of 58 fractions were collected. They were combined into 20 fractions and lyophilized.

#### LC MS/MS

The fractionated phosphopeptides were analysed by LC MSMS using a fully automated Ultimate 3000 RSLC nano System (Thermo Scientific) fitted with a 100 µm x 2 cm PepMap100 C18 nano trap column and a 75 µm × 25 cm reverse phase C18 nano column (Aclaim PepMap, Thermo Scientific). Lyophilised phosphopeptides were dissolved in solvent A (2% MeCN, 0.1% formic acid) and eluted with a linear gradient from 4 to 50% solvent B (80% MeCN, 0.1% formic acid) over 110 min with a flow rate of 300 nl/min. The outlet of the nano column was directly interfaced via a nanospray ion source to a Q Exactive Plus mass spectrometer (Thermo Scientific). The mass spectrometer was operated in standard data dependent mode, performing a MS full-scan in the m/z range of 350-1600, with a resolution of 140000. This was followed by MS2 acquisitions of the 15 most intense ions with a resolution of 35000 and NCE of 33%. MS target values of 3e6 and MS2 target values of 1e5 were used. The isolation window of precursor was set at 1.2 Da and sequenced peptides were excluded for 40 sec.

#### Data analysis

The acquired MSMS raw files from the analysis above were processed using Proteome Discoverer (version 2.1, Thermo Scientific). MSMS spectra were searched against the January 2016 *Dictyostelium* discoideum UniProt FASTA database, using Mascot (version 2.4, Matrix Science) search engine. Cysteine carbamidomethylation was set as a fixed modification, while methionine oxidation, N-terminal acetylation (protein), phosphorylation (STY) and TMT6plex (peptide N-terminus and Lysine) were selected as variable modifications. Other parameters were set to default values. The abundance values of TMT reporter ions were normalized to total peptide amount. Two output files were generated; one listing the proteins identified and one for peptides. Only peptides that were identified with a high confidence (False discovery rate-FDR-of <1%) based on a target decoy approach were included in these results. The protein output file was filtered to only contain proteins which passed the threshold FDR of 1% and exported as an excel file. The peptides table was filtered for phosphopeptides with quantitative values and exported as an excel file.

The phosphopeptides were filtered by reference to the fluid uptake data obtained earlier: ≥5-fold decreased in the PkbA-/PkbR1-mutant compared to Ax2, ≥2-fold decreased in the PkbR1-+ LY294002 cells and have at least a 20% decrease in one of the PkbA-and PkbR1-samples. Only phosphopeptides meeting these criteria in at least 2 of the 3 repeats were accepted. The likely kinase responsible for the phosphorylation was then predicted using NetPhos 3.1.

### Quantification and statistical analysis

Statistical analysis was performed using Graphpad prism software, which was used to generate the presented graphs. When there were more than two samples to compare, one-way anova was performed before any further statistical tests. Unless otherwise specified an unpaired t-test was performed and p values ≤0.01 are marked with **, those ≤0.05 * and those ≤0.1 are indicated in text. Unless otherwise stated the mean is plotted with the error bars showing the SEM.

## Acknowledgements

The authors thank the Kay lab for their input into this project and the LMB microscopy and flow cytometry facilities for excellent scientific and technical support. We are also very grateful to Douwe Veltman for modifying his membrane finder software, and Peter Devreotes for providing the PDK mutants.

## Data availability

The phosphoproteomic data set generated is available in Data S1.

